# Your place or mine? The neural dynamics of personally familiar scene recognition suggests category independent familiarity encoding

**DOI:** 10.1101/2023.06.29.547012

**Authors:** Hannah Klink, Daniel Kaiser, Rico Stecher, Géza Gergely Ambrus, Gyula Kovács

**Author notes:** These authors contributed equally as senior authors (Géza Gergely Ambrus, Gyula Kovács). HK: Institute of Neurology, Universitätsklinikum Jena, D-07747 Jena, Germany. DK: Mathematical Institute, Justus-Liebig-University Gießen, D-35392 Gießen, Germany. RS:Mathematical Institute, Justus-Liebig-University Gießen, D-35392 Gießen, Germany. GK: Institute of Psychology, Friedrich Schiller University Jena, D-07743 Jena, Germany. GGA: Department of Psychology, Bournemouth University, Poole House, Talbot Campus, Fern Barrow, Poole, Dorset, BH12 5BB, United Kingdom.

## Abstract

Recognizing a stimulus as familiar is an important capacity in our everyday life. Recent investigation of visual processes has led to important insights into the nature of the neural representations of familiarity for human faces. Still, little is known about how familiarity affects the neural dynamics of non-face stimulus processing. Here we report the results of an EEG study, examining the representational dynamics of personally familiar scenes. Participants viewed highly variable images of their own apartments and unfamiliar ones, as well as personally familiar and unfamiliar faces. Multivariate pattern analyses were used to examine the time course of differential processing of familiar and unfamiliar stimuli. Time-resolved classification revealed that familiarity is decodable from the EEG data similarly for scenes and faces. The temporal dynamics showed delayed onsets and peaks for scenes as compared to faces. Familiarity information, starting at 200 ms, generalized across stimulus categories and led to a robust familiarity effect. In addition, familiarity enhanced category representations in early (250 – 300 ms) and later (>400 ms) processing stages. Our results extend previous face familiarity results to another stimulus category and suggest that familiarity as a construct can be understood as a general, stimulus-independent processing step during recognition.

**Highlights:**

1. Whether a face or scene is familiar can be decoded from the EEG signal with very similar temporal dynamics, starting at 200 ms and peaking around 400 ms after stimulus onset.
2. The neural dynamics of this familiarity information generalizes across stimulus categories.
3. Familiarity modulates stimulus category representations from 200 ms after stimulus onset, indicating deeper processing of familiar as compared to unfamiliar stimuli already during early processing stages.

## Introduction

Stimulus recognition is a multifaceted task that requires several processing steps, among which the construction of familiarity– the feeling that a stimulus has been encountered before (Rugg and Yonelinas 2003) – stands out as a critical component. For face stimuli, research over the past few decades, has demonstrated a significant behavioral advantage for familiar faces, including fast and accurate recognition (Burton, 2005), automatic processing (Yan et al., 2017), and effortless matching of the same familiar identity across highly variable features (Jenkins et al. 2011). Neuroimaging and electrophysiological studies have recently expanded upon existing behavioral data to explore the neural representation of face familiarity. Functional magnetic resonance imaging (fMRI) studies have identified several cortical areas involved in processing face familiarity, which has facilitated the development of multiple models for understanding face familiarity (Duchaine and Yovel 2015; Kovács 2020). Given its low temporal resolution, fMRI is not well-suited for capturing the temporal dynamics of these processes, therefore researchers have utilized uni- and multivariate EEG/MEG analyses to study the temporal aspects of face familiarity processing. Univariate event-related potential (ERP) studies have identified various face-specific ERP components that are sensitive to familiarity. Regarding the first face-selective ERP component (the N170; Bentin et al., 1996), there is currently no consensus on its sensitivity to face familiarity, as conflicting findings have been reported (Gosling and Eimer 2011; Barragan-Jason et al. 2015; Andrews et al. 2017). It is possible that the sensitivity of the N170 to face familiarity is contingent upon task context and may be more pronounced when familiarity is especially high (Johnston et al. 2016; Caharel and Rossion 2021). This interpretation is consistent with findings supporting the early, feedforward modulation of perceptual information by high familiarity (Karimi-Rouzbahani et al. 2021). In contrast to the N170, the subsequent N250 ERP component is clearly modulated by familiarity, showing a more pronounced negative deflection for familiar compared to unfamiliar faces (Andrews et al. 2017). The N250 typically emerges between 200 and 400 ms after stimulus onset and has an occipitotemporal scalp distribution. It is assumed to be related to the comparison of perceptual inputs to stored face representations (Wiese et al. 2019). Huang and colleagues (Huang et al. 2017) found a correlation between the N250 and reaction times in a face matching task, thus demonstrating the direct behavioral relevance of this component. Recent studies using multivariate pattern analysis (MVPA) have confirmed these results, demonstrating that information regarding face familiarity can be detected in the signal as early as 200 ms after stimulus onset (Ambrus et al. 2021; Dalski, Kovács, and Ambrus 2022; Li et al. 2022), with some studies suggesting an even earlier emergence of familiarity effects (Bayer et al. 2021) or an early modulation of stimulus properties by familiarity (Dobs et al. 2019).

Although the exact timing of the onset of familiarity information remains a topic of ongoing research, both ERP and MVPA studies consistently indicate that the most robust signal for face familiarity is observed at around 400 ms (Ambrus et al. 2021). This time-frame around 400 ms overlaps strongly with a recently identified ERP component labelled as the ‘Sustained Familiarity Effect’ (SFE; Wiese et al., 2019). The SFE is a strong and reliable indicator of facial familiarity that develops only after a certain level of exposure to an individual and it is theorized to reflect the accumulated mnemonic, social, and affective information (Popova and Wiese 2022, 2023). Thus, the SFE may represent the integration of perceptual and stored representations, enabling the recognition of familiar individuals. Although at least a partial functional separation between the N250 and SFE has been proposed, recent MVPA studies suggest that there is a continuity in the signal, indicating a single, long-lasting and robust neural process underlying face familiarity that generalizes across participants, and even experiments (Dalski, Kovács, and Ambrus 2022; Dalski, Kovács, Wiese, et al. 2022).

While recognition memory, including familiarity and recollection (Rugg and Yonelinas 2003; Dimsdale-Zucker et al. 2022) has been explored for various types of stimuli, there has been limited research on whether the underlying neural processes of familiarity differ or are similar across stimuli categories, and if they generalize across different types of stimuli (Kwon et al. 2022). Recently, Ambrus (2022) conducted a cross-experiment classification analysis based on data from prior studies and found a significant overlap in neural signals of recognition memory processes between 400-600 ms, regardless of sensory modality, stimulus type, or memory age. Although these findings provided evidence for shared recognition effects across different types of stimuli, a systematic, direct comparison of the temporal dynamics of memory processes for face and non-face stimuli is still lacking.

Therefore, in the current study we investigated the generalizability of the familiarity signal between faces and another stimulus category, scenes. Despite the available evidence from fMRI studies on the differential response of brain regions to familiar and unfamiliar scenes (Epstein, Higgins, et al. 2007; Epstein, Parker, et al. 2007; Bainbridge and Baker 2022), there is a paucity of research investigating the temporal dynamics of scene recognition, especially in comparison to the extensive M/EEG literature on face familiarity. In a recent cued recall experiment, Treder and colleagues (2021) used learned object/scene and scene/object associations to explore the time course of the switch from perceiving the environment to retrieving goal-relevant memories, but, to the best of our knowledge so far, no study contrasted familiar and unfamiliar scene processing directly using M/EEG. Thus, the first aim of the current study is to fill this gap by examining the temporal dynamics of personally familiar and unfamiliar scene recognition. We selected personally familiar scenes, specifically images of the participants’ apartments as prior studies have shown that personal familiarity tends to elicit the strongest recognition signals for face stimuli (Ambrus et al. 2021; Dalski, Kovács, and Ambrus 2022; Li et al. 2022; Popova and Wiese 2022). In addition to personally familiar scenes, we also presented personally familiar and unfamiliar faces to compare the emerging scene familiarity signals to the previously established face familiarity representations. The second aim of the study was to test for a common neural signature of familiarity across faces and places, probing the generalizability of the familiarity signal between these two stimulus categories.

## Methods and Materials

### Participants

32 participants (7 male; average age: 23.13, SD = 2.96) took part in the study in exchange for monetary compensation or partial course credits. Data from 3 participants (2 male) were excluded from the final analysis due to excessive noise in the EEG. Another 2 participants did not give consent to their EEG data being published. The final sample includes data from 27 participants (5 male; average age: 23.04, SD = 2.91). This sample size is comparable to that of previous studies investigating the neural correlates of personal face familiarity (Ambrus et al., 2021; Bayer et al., 2021; Li et al., 2022; Wiese et al., 2019). Participants reported no history of neurological conditions, had normal or corrected-to-normal vision, and were all right-handed. They were recruited through the institute’s mailing list or personal contacts. Participants provided written and informed consent prior to participating. The study was conducted in accordance with the guidelines outlined in the Declaration of Helsinki and was approved by the ethics committee of the Friedrich Schiller University Jena.

### Stimuli

The stimuli were images of faces and scenes. Participants provided 20 photographs of faces of personally familiar people (4 images of 4 identities each + 4 images of the participants’ own face) and 20 photographs of their own apartment (Figure 1). Personally familiar identities were defined as “people you are very close to”. Photographs of apartments did not feature people or animals and depicted typical interior scenes such as a home-office desk in front of a window or a bathroom sink with several objects scattered around. The set of unfamiliar faces and scenes comprised 4 images per 5 identities, and 20 photographs of the apartments, respectively. These were provided by another randomly selected participant. We ensured that the participants did not know each other and never visited the other participant’s apartment. To further expand the pool of unfamiliar stimuli, we included images provided by student assistants, colleagues, and the authors’ families. After the EEG recording, participants were asked to fill out a questionnaire to control for accidental familiarity. They were asked to rate their level of familiarity with the unfamiliar stimuli on a scale ranging from 0 to 5, with 0 indicating complete unfamiliarity and 5 indicating high personal familiarity. Note that we do not analyze the own-face trials in the present study as these results will be published in a separate report.

**Figure 1.**
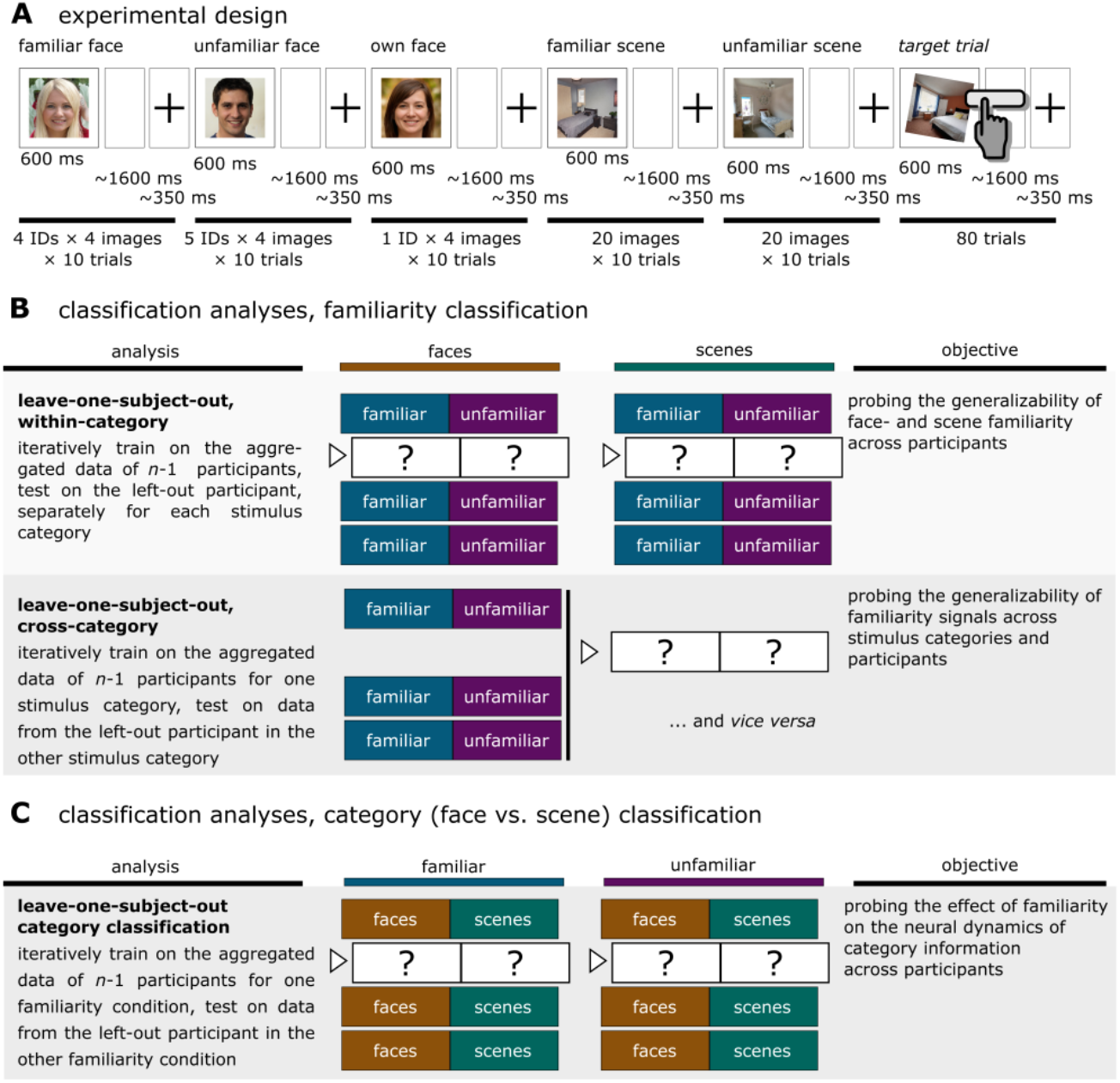
Experimental design and classification pipelines. **(A) Stimulus presentation**. Personally familiar and unfamiliar faces, the participants’ own face, and familiar and unfamiliar scenes were presented in a pseudorandom order. The presentation time was 600 ms, with a ca. 1650 ms inter-stimulus interval (fixation cross: 350 ms ± 100 ms temporal jitter, blank screen 1650 ms ± 200 ms temporal jitter). To maintain attention, participants were instructed to press the space bar for images rotated by 10° (target trials, not included in the analyses); no other response was required. **(B) Familiarity classification**. To examine the temporal dynamics of face and scenes processing, as well as the generalizability of the familiarity signal, leave-one-subject-out familiarity decoding was performed both within-category (trained on faces / scenes, tested on faces / scenes) and cross-category (face and scene stimuli as test categories each). **(C) Category classification**. To examine the effect of familiarity on stimulus category processing, leave-one-subject-out category decoding was performed with familiar and unfamiliar stimuli as test categories each. Example faces were generated by AI for illustration purposes (https://thispersondoesnotexist.com/).

### Experimental design

In total, 800 trials (80 stimulus images each repeated 10 times) were presented during the EEG recording. Stimuli were displayed on a grey background (hex color format: #808080; see **Figure 1A** for trial structure) for 600 ms, followed by gap of 1600 ms. All stimuli subtended 11.42° of visual angle. The image sequence was pseudo-randomized in a way that two images of the same category (e.g., the same face identity) could not appear back-to-back. To maintain attention throughout the experiment, 10% of the trials were target trials, where images were rotated 10° clockwise. The participants were instructed to press a button when these stimuli appeared. The experiment was written in PsychoPy v2021.2.3 (Peirce et al. 2019).

### EEG acquisition and preprocessing

EEG was recorded using a 64-channel Biosemi Active II system (512 Hz sampling rate) in a dimly lit, electrically shielded, and sound-attenuated chamber. EEG data preprocessing was carried out using MNE-Python (Gramfort et al. 2013). Data were bandpass filtered between 0.1 and 40 Hz, segmented between −200 and 1200 ms and baseline-corrected to the 200 ms preceding stimulus presentation. The data were downsampled to 100 Hz (resulting in 140 time points) and no artifact rejection was performed (Grootswagers et al. 2017; Delorme 2022).

### Analysis pipelines

Within- and cross-participant classification analyses were conducted using linear discriminant analysis (LDA) classifiers. Training data trial counts were always balanced on the participant level by under-sampling to the minimum image and trial count in the classes of interest. Time-resolved classification (Grootswagers et al. 2017), spatio-temporal searchlight, and temporal generalization (King and Dehaene 2014) analyses were conducted. In the time-resolved classification procedure, the classifiers were trained and tested at each of the 140 time points to distinguish between the classes of interest: familiar versus unfamiliar stimuli in the familiarity classification analysis or faces versus scenes in the category classification analysis. This procedure was used to assess the decoding accuracy for individual participant data in the within-participant analyses, and for data from a participant not included in the training set in the leave-one-participant-out analyses. Temporal generalization followed a similar approach, where classifiers trained at a specific time point were used to test data from all other time points, resulting in a cross-temporal classification accuracy matrix. Time-resolved classification and temporal generalization was performed over all electrodes as well as pre-defined regions of interest, which included six scalp locations along the median (left and right) and coronal (anterior, center, and posterior) planes. The spatio-temporal searchlight procedure involved systematically testing each channel by training and testing on data from that channel and its neighboring electrodes, employing the same time-resolved analysis as described previously. For further details, see **Supplementary Information 1**.

### Familiarity classification

Cross-participant classification allows for better testing of generalization as it assesses the ability to classify stimuli across individuals, ensuring that the findings are not specific to individual characteristics. Furthermore, using participant-unique stimulus sets helps reduce stimulus-specific effects such as low-level image properties, resulting in increased generalizability of the findings. Previous studies investigating familiarity representations have consistently demonstrated strong generalization across participants within the same stimulus category (see e.g., Dalski, Kovács, and Ambrus 2022; Li et al., 2022).

As the major aim of the current study was to test the generalization of familiarity effects across stimulus categories, here we only present the results of the cross-participant analysis in detail. For an in-depth description of training and test sets, as well as for the results of the within-subject classification analyses see **Supplementary Information (https://osf.io/m9q74/)**.

To characterize the temporal dynamics of the generalizability of the familiarity signals across participants, leave-one-subject-out classification analyses were performed. We used a leave-one-out cross-validation approach, in which we held out the data from one participant for testing and used the aggregated data from all the other participants for training. Image trial counts for scenes, and image and identity trial counts for faces, were equalized for each participant to ensure a fully balanced dataset. A moving average of 30 ms (3 consecutive time points) was applied to all classification accuracy data at the participant level (Ambrus et al. 2021; Dalski, Kovács, and Ambrus 2022)

Familiarity classification was conducted both within and across stimulus categories. Within-category classification refers to using data from the same stimulus type for both training and testing (e.g., training the classifier on faces and subsequently testing on faces, as well as training and testing on scenes). This was done to examine the temporal evolution of the familiarity signals for each stimulus type separately. Cross-category classification, on the other hand, refers to using the data from one stimulus type for training and the data from the other stimulus type for testing (e.g., training the classifier on faces and testing it on scenes, or vice versa). This analysis allows us to identify the shared neural signals of familiarity that are present across stimulus types and across participants (leave-one-participant-out analyses).

### Category classification

To test if familiarity also modulates the representational dynamics of category information, an additional cross-participant face versus scene classification was performed separately on evoked responses for familiar and for unfamiliar trials.

### Statistical inference

In time-resolved analyses, classification accuracies were entered into two-tailed, one-sample cluster permutation tests (10,000 iterations) against chance (50%). In temporal generalization and searchlight analyses, two-tailed spatio-temporal cluster permutation tests were used against chance level (50%), with 10,000 iterations. Statistical analyses were conducted using python, MNE-Python (Gramfort et al. 2014), scikit-learn (Pedregosa et al. 2011), and SciPy (Virtanen et al. 2020).

## Results

### Familiarity decoding of scene and face stimuli

**Figure 2A** presents the results of the familiarity classification analysis, when performed within stimulus categories. For faces, we found significant clusters of familiarity information (cluster *p*s < 0.0001) at 180 ms, with a peak at 430 ms and lasting until the end of the epoch (peak Cohen’s *d* = 1.8). For scenes, familiarity information emerged slightly later at 210 ms, peaked at 380 ms and lasted until 1030 ms (peak Cohen’s *d* = 1.4). A significant difference between the two stimulus categories was only observed in an early time-window between 120 and 290 ms, with a maximum difference at 240 ms (cluster *p* = 0.008, peak Cohen’s *d* = 0.73) with faces leading to higher classification accuracies than scenes. The spatio-temporal searchlight analyses revealed similar patterns. Robust clusters (cluster *p*s < 0.0001), encompassing all electrodes were observed for faces (onset: 120 ms, peak at 430 ms over PO10, peak Cohen’s *d* = 2.2) as well as for scenes (onset: 190 ms, peak at 1190 ms over PO10, peak Cohen’s *d* = 1.6). Temporal generalization results (**Figure 3A-B**) yielded robust (cluster *p*s < 0.0001) and sustained clusters until the end of the epoch both in the case of faces (from 110 ms train time and 120 ms test time, peak Cohen’s d = 1.859) and scenes (from 170 ms train time and 180 ms test time, peak Cohen’s *d* = 1.486).

**Figure 2.**
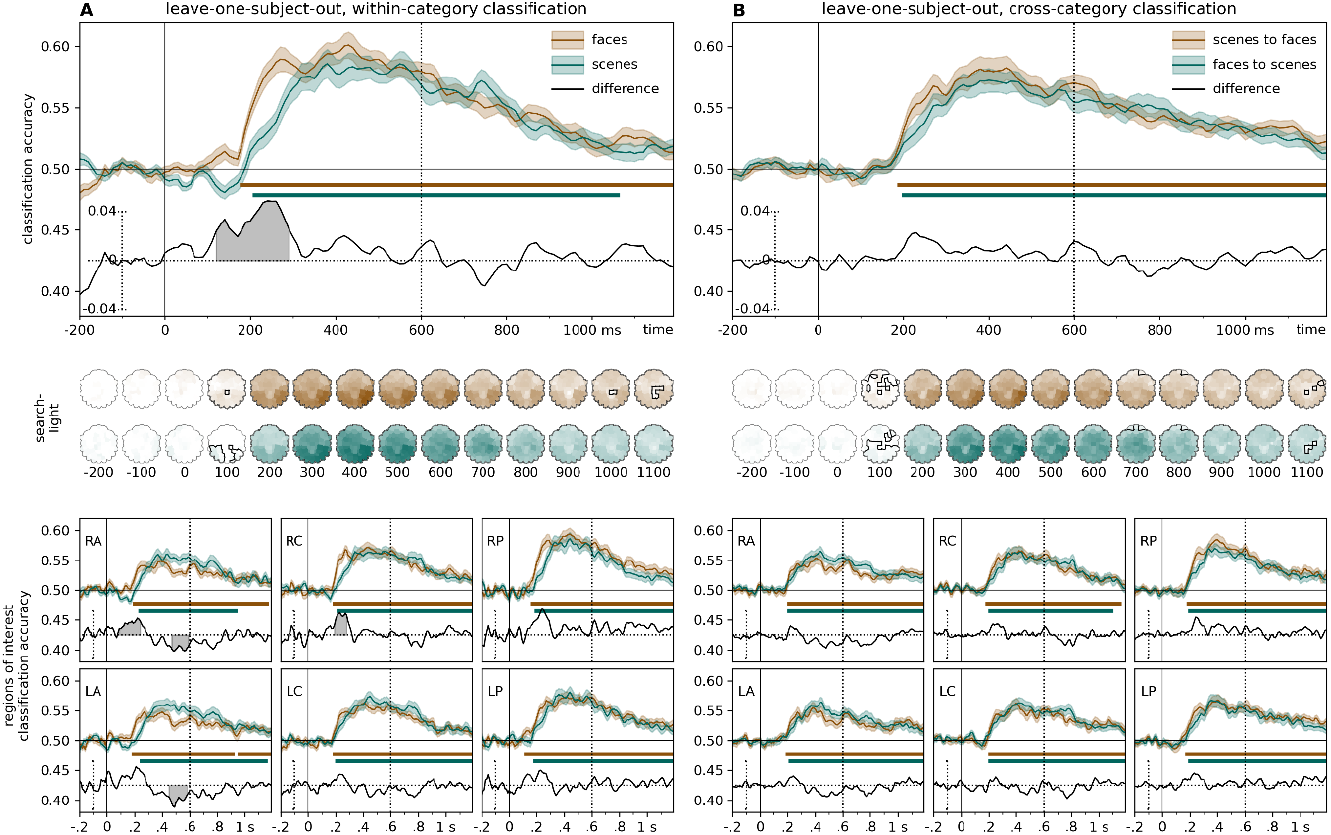
Time-resolved, leave-one-subject-out classification of familiarity. Classifiers were trained to categorize ERPs for familiar and unfamiliar stimuli. **(A) Within-category classification**. Training and testing for familiarity was performed within the same stimulus category (*face*: iteratively trained on data form *n*-1 participants’ ERPs for faces, tested on one participants’ ERPs for faces. *scene*: iteratively trained on data from *n*-1 participants’ ERPs for scenes, tested on one participants’ ERPs for scenes). **(B) Cross-category classification**. Training and testing for familiarity was performed on different stimulus categories (*face*: iteratively trained on *n*-1 participants’ ERPs for scenes, tested on one participants’ ERPs for faces. *scene*: iteratively trained on *n*-1 participants’ ERPs for faces, tested on one participants’ ERPs for scenes). Two-sided cluster permutation tests, *p* < .05. Top panels show results for analyses on all electrodes. Middle Panels: spatio-temporal searchlight results are shown as scalp maps, with classification accuracy scores averaged in 100 ms steps. Sensors and time points belonging to the significant cluster when tested on faces are shown in the top row, sensors and time points belonging to the significant cluster when tested on scenes are shown in the bottom row. (Two-sided spatio-temporal cluster permutation tests, *p* < .05). Bottom panels: ROI analyses. The same analysis as in the top panel, but repeated for six rep-defined electrode clusters separately(Ambrus et al. 2019). RA/LA: right/left anterior, RC/LC: right/left central, RP/LP: right/left posterior. The vertical line at 600 ms denotes the end of the stimulus presentation. For detailed statistics, see **Supplementary Table 1J-O**., and **Supplementary Table 3G-J**.

**Figure 3.**
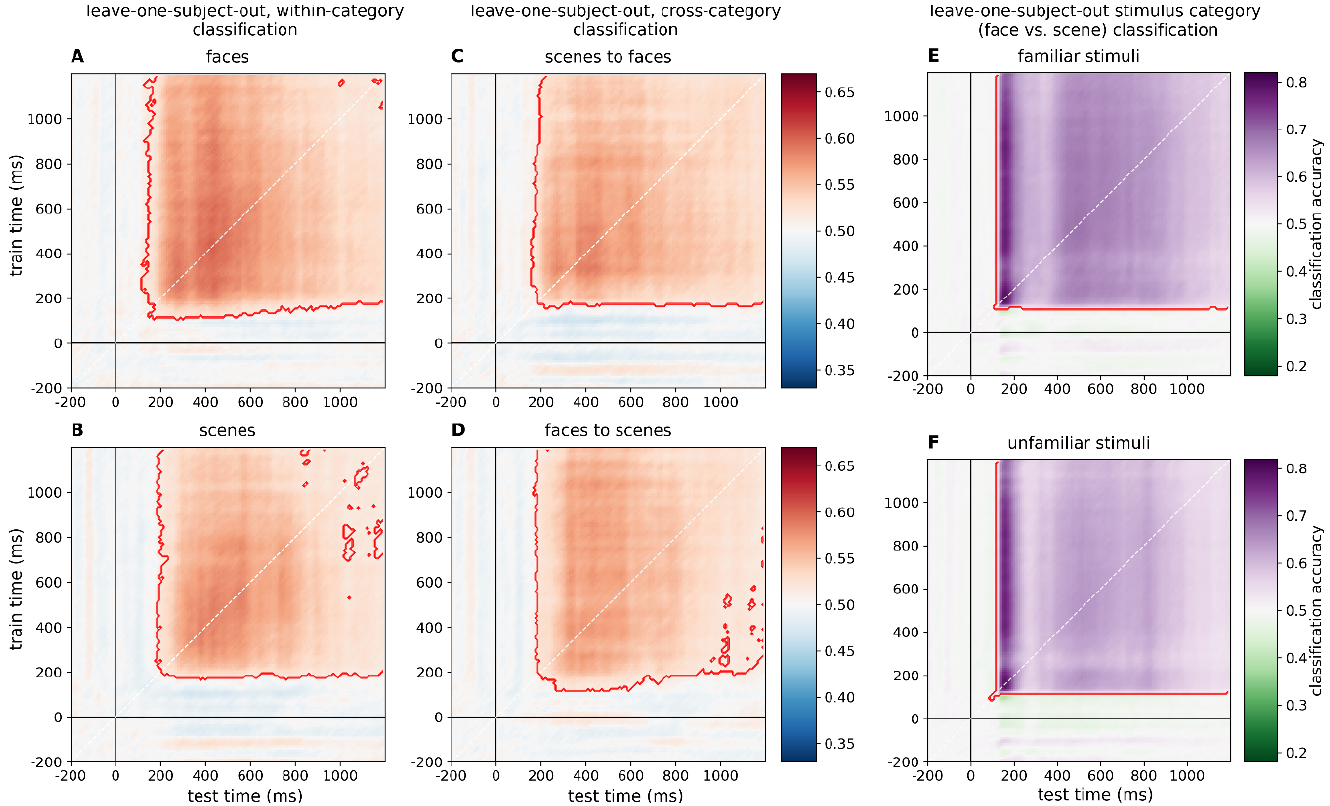
**Temporal generalization analyses** of familiarity and category (leave-one-subject-out classification). **(A, B) Within-category classification of familiarity**. Classifiers were trained to categorize ERPs for familiar and unfamiliar stimuli. (*face*: iteratively trained on data form *n*-1 participants’ ERPs for faces, tested on one participants’ ERPs for faces. *scene*: iteratively trained on data form *n*-1 participants’ ERPs for scenes, tested on one participants’ ERPs for scenes). **(C, D) Cross-category classification of familiarity**. (*face*: iteratively trained on *n*-1 participants’ ERPs for scenes, tested on one participants’ ERPs for faces. *scene*: iteratively trained on *n*-1 participants’ ERPs for faces, tested on one participants’ ERPs for scenes). **(E, F) Category classification**. Stimulus category was classified for familiar and unfamiliar trials separately and iteratively trained on *n*-1 participants’ ERPs for the two stimulus categories (faces and scenes), and tested on a left-out participants’ ERPs. Two-sided cluster permutation tests, *p* < .05. For the results of the temporal generalization analyses in the pre-defined regions of interest, see **Supplementary Table 2**.

Within-participant familiarity decoding yielded similar results as the above described cross-participant analysis. Onset, peak values, and corresponding statistics can be found in **Supplementary Table 1A-C** and **Supplementary Table 2A-B**. For the details of the searchlight results, see also **Supplementary Table 3**.

### Familiarity decoding across stimulus categories

Although the previous analysis reveals that the neural patterns of familiarity representation exhibit remarkable similarity for both scenes and faces, it remains a possibility that these findings are attributed to two distinct or partially distinct neural mechanisms that share similar dynamics. To confirm whether the neural representations of face and scene familiarity are supported by common mechanisms, we conducted cross-category classification by training the classifier on data from one category and testing it on data from the other category (**Figure 2B**).

In this analysis, we also found significant clusters lasting for a prolonged period (cluster *p*s < 0.0001) in both decoding directions. Cross-category classification was found for scenes to faces from 190 ms with a peak at 440 ms (peak Cohen’s *d* = 1.6) as well as for faces to scenes from 200 ms with a peak at 400 ms (peak Cohen’s d = 1.5). A sharp rise in familiarity information was found at around 200 ms after stimulus onset, followed by a plateau until 450 ms with a peak at approximately 400 ms, after which a slow and steady decrease was observed. General familiarity information persisted until the end of the epoch, reflecting sustained processing. Again, spatio-temporal searchlight analyses revealed similar patterns for both decoding directions. Robust clusters (cluster *p*s < 0.0001), encompassing all electrodes, were observed for scenes to faces (onset: 160 ms, peak at 440 ms over TP8, peak Cohen’s *d* = 2.0) as well as faces to scenes (onset: 180 ms, peak at 430 ms over TP8, peak Cohen’s *d* = 1.8). Temporal generalization (**Figure 3C-D**) yielded robust (cluster *p <* 0.0001) and sustained clusters, for scenes to faces (train-time onset: 160 ms, test-time onset: 160 ms, peak Cohen’s *d* = 1.2) as well as for faces to scenes (train-time onset: 120 ms, test-time onset: 170 ms, peak Cohen’s *d* = 1.6).

For the results of the cross-category analyses in the six pre-defined regions of interest electrode clusters, see **Supplementary Table 1J-O** and **Supplementary Table 2G-J**. For the details of the searchlight results, see **Supplementary Table 3G-J**. When performed within-participant, cross-category decoding yielded similar results as the above described cross-participant, cross-category familiarity decoding. Onset, peak values, and corresponding statistics can be found in **Supplementary Table 1D-F** and **Supplementary Table 2C-D**. For the details of the searchlight results, see **Supplementary Table 3**.

### The effect of familiarity on category representation

In our previous analyses, we demonstrated a differential representation for familiar and unfamiliar scenes and faces. Next, we investigated whether familiarity also improved the neural discriminability between the categories. Specifically, we examined whether classifiers were more successful in distinguishing between faces and scenes when they were familiar as opposed to unfamiliar. Time-resolved leave-one-participant-out classification of stimulus category (**Figure 4**, top panel) for both familiar and unfamiliar stimuli revealed robust (cluster *p*s < 0.0001) and sustained clusters with an early onset. For familiar stimuli, the cluster onset was at 110 ms, with a peak at 160 ms (peak Cohen’s *d* = 3.8), while for unfamiliar stimuli, the onset of the significant cluster was at 90 ms, with a peak at 160 ms as well (peak Cohen’s *d* = 3.8). Importantly, both early and late differences between familiar and unfamiliar stimuli were observed. A cluster in a brief and early time-window, between 220 and 300 ms (peak at 240 ms, cluster *p* = 0.040, peak Cohen’s *d* = 0.69), and two late clusters (between 400 and 800 ms, peak at 450 ms, cluster *p* = 0.0005, peak Cohen’s *d* = 1.0, and from 830 ms, peak at 1010 ms, cluster *p* = 0.0011, peak Cohen’s *d* = 0.79) were significant, indicating significantly higher category classification performance for familiar stimuli than for unfamiliar stimuli. Searchlight analyses (**Figure 4**, middle panel) yielded strong category decoding for both familiar and unfamiliar stimuli, both peaking over the PO8 electrode (cluster *p*s < 0.0001), with a cluster onset at 70 ms for familiar (peak Cohen’s *d* = 4.4), and a cluster onset at 60 ms for unfamiliar (peak Cohen’s *d* = 4.0) stimuli. In both cases, temporal generalization (**Figure 3E-F**) yielded robust (cluster *p <* 0.0001) and sustained clusters, both for familiar stimuli (train-time onset: 110 ms, test-time onset: 110 ms, peak Cohen’s *d* = 3.7) and for unfamiliar stimuli (train-time onset: 90 ms, test-time onset: 90 ms, peak Cohen’s *d* = 3.8).

**Figure 4.**
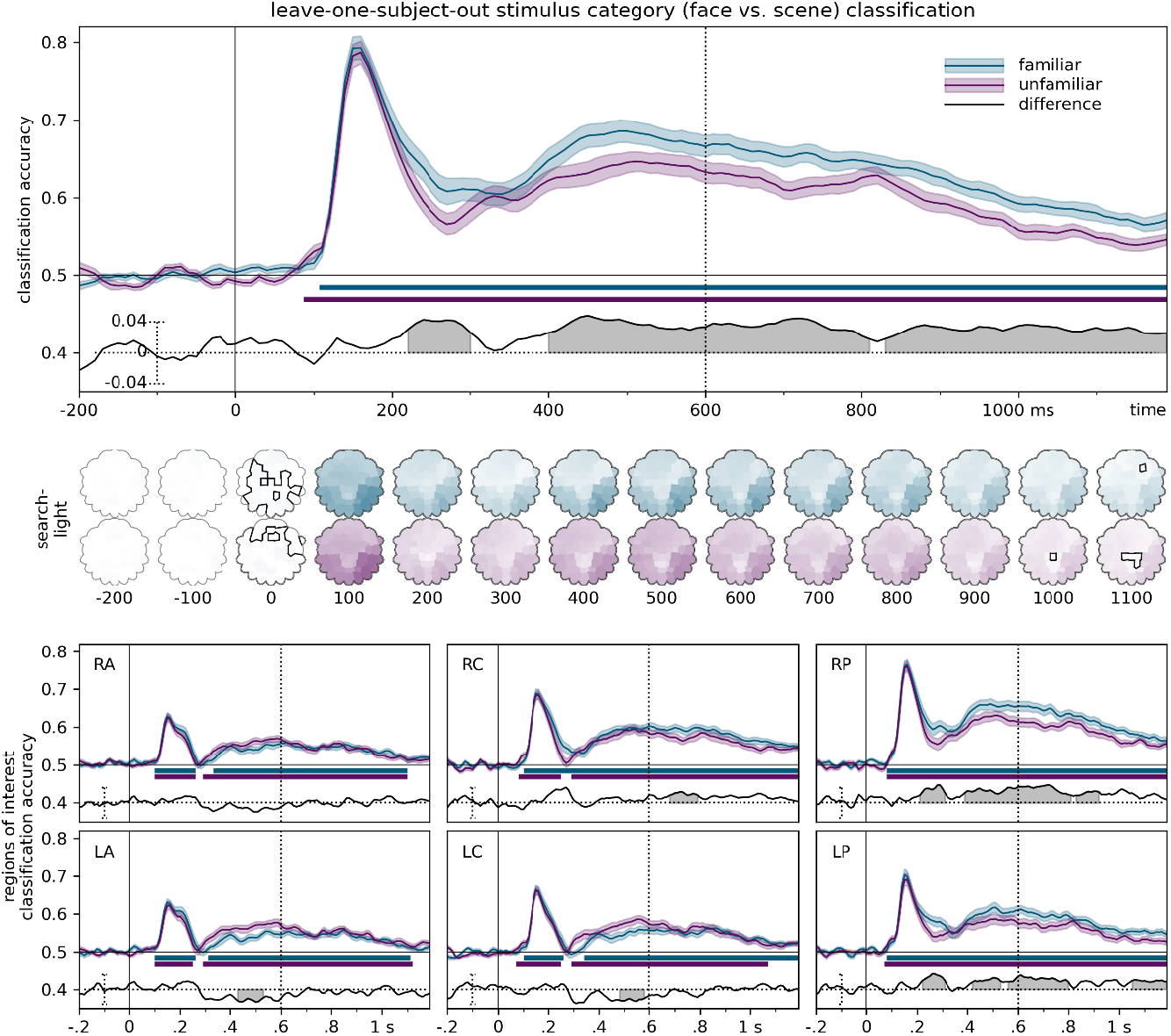
Time-resolved, leave-one-participant-out classification of stimulus category. Classifiers were trained, across participants, to categorize stimulus category (faces vs. scene), for familiar and unfamiliar trials separately. Two-sided cluster permutation tests, *p* < .05. Spatio-temporal searchlight results are shown as scalp maps, with classification accuracy scores averaged in 100 ms steps. Sensors and time points belonging to the significant cluster when tested on familiar stimuli are shown in the top row, sensors and time points belonging to the significant cluster when tested on unfamiliar stimuli are shown in the bottom row. (Two-sided spatio-temporal cluster permutation tests, *p* < 0.05). RA/LA: right/left anterior, RC/LC: right/left central, RP/LP: right/left posterior. The vertical line at 600 ms denotes the end of the stimulus presentation. For detailed statistics, see **Supplementary Table 1P-R**., and **Supplementary Table 3K-L**.

Results of the leave-one-subject-out stimulus category time-resolved cross-classification and temporal generalization analyses for the pre-defined regions of interest can be found in **Supplementary Table 1P-R** and **Supplementary Table 2K-L**. For the details of the searchlight results, see **Supplementary Table 3L-L**. When performed within-participant, the category decoding yielded similar results as the above described cross-participant category decoding. Onset, peak values, and corresponding statistics can be found in **Supplementary Table 1G-I** and **Supplementary Table 2E-F**. For the details of the searchlight results, see **Supplementary Table 3E-F**.

## Discussion

We investigated the temporal dynamics of familiarity processing for personally familiar scenes and explored the generalization of familiarity information across faces and scenes. The key results of the current study are as follows: 1) The temporal dynamics of familiarity processing for personally familiar scenes resembles that of personally familiar faces. A notable difference between the two categories is the onset of familiarity information, which is delayed for scenes. 2) Familiarity information emerges at 200 ms, generalizes between two personally familiar categories and leads to a robust and sustained familiarity effect that is independent of stimulus category. 3) Familiarity also modulates stimulus category representations by enhancing it in both early (250 – 300 ms) and later (>400 ms) processing stages.

### Similar temporal dynamics of familiarity processing for face and non-face stimuli

Familiarity, reflecting previous exposure to a particular stimulus, serves as the basis of correct recognition memory (Yonelinas, 2002), and familiarity has long been a topic of interest in face perception research. Multiple recent human electrophysiological studies have characterized the processing dynamics of face stimuli, and observed robust and long-lasting familiarity effects (Wiese et al. 2019; Ambrus et al. 2021; Dalski, Kovács, and Ambrus 2022). However, due to the absence of empirical studies directly comparing this familiarity effect between faces and other highly familiar non-face stimuli, it remains still an open question if the observed familiarity effect is specific to the stimulus category of faces. We propose it to be a general signature of familiarity per se, representing a critical stage in the process of recognition memory that is not specific to any particular stimulus category. When exploring familiarity across faces and non-face stimulus categories, a critical challenge is that in our every-day life we tend to have extensive exposure to the faces we know (Young & Burton, 2018). As a result, our ability to recognize highly familiar faces, such as those of our family members, is superior to the recognition of other faces and other, non-face objects (Contini et al. 2017; Ramon and Gobbini 2018). Consequently, it is difficult to find non-face stimulus categories where exemplars are similarly well known to the observer as personally familiar faces. The major strength of this current study is that we used images of our participants’ own apartments as personally familiar scenes. These scenes provided a source of stimuli for which participants had acquired extensive prior exposure; thus we were able to compare the neural dynamics of personally and highly familiar face and non-face stimuli in a within-subject design. We hypothesized that if the previously observed familiarity effect in the neural signal is indeed a general signature of familiarity, we should find very similar temporal dynamics for both stimulus categories.

Indeed, the neural patterns reflecting familiarity generalized well between faces and scenes, similarly in both decoding directions. Furthermore, cross-category decoding also led to significant classification performance across participants, indicating its independence of both stimuli and participants. Our findings suggest that personally familiar faces and scenes are processed in sufficient detail to elicit a stimulus-independent and robust familiarity signal by 200 ms post stimulus onset. This effect had previously been shown for faces across participants, experiments, and different qualities of familiarity (Dalski, Kovács, and Ambrus 2022). Here we present evidence that a similar effect is present in across participants for personally familiar scenes. It is thus very likely that this effect extends to other experimental paradigms, different qualities of familiarity, and even different stimulus categories. Indeed, a recent cross-category classification study by Ambrus (2022) showed that the same processing dynamics observed for familiar faces also apply to other categories and experimental paradigms, such as familiar and remembered objects, remembered (vs. forgotten) object-scene associations, and even to music, reflecting a generalization across sensory modalities. Familiarity effects were observed for all categories examined in the analysis at a relatively later time range (post 300-400 ms), while a shared and sustained effect was observed earlier (at 200 ms) only for personally familiar faces and experimentally familiarized objects later judged as subjectively familiar, but not for those judged as remembered. This finding of an earlier effect for objects aligns with the current study’s results, further strengthening the conclusion that the observed familiarity effect is not specific to faces. Instead, all evidence indicates that familiarity, as previously established, reflects a general and stimulus-independent process triggered by previously encountered exemplars of an object category, which can be seen as early as 200 ms after stimulus onset. It is likely that this process is related not just to a familiarity judgement per se, but, especially at later stages, to specific recall processes involving episodic memory and recollection (Ambrus 2022).

### Dynamics of familiar scene processing

The fact that familiarity information can be generalized across different stimulus categories suggests that it is processed by the same mechanisms and neural signals. Although the trajectory of face familiarity information has previously been established (e.g., Ambrus et al. 2021), empirical studies exploring the generalizability of familiarity information across different stimulus types remain scarce. Here, through direct investigation of cross-participant familiarity effects for both faces and scenes we provide compelling evidence in support of the generalizability of familiarity information, suggesting that the classification of familiarity is not restricted to particular stimulus categories or individuals. Our findings are in alignment with the previously established temporal patterns of familiarity processing in studies using event-related potentials and multivariate pattern analysis with face stimuli. Critically, the scene familiarity dynamics of the current study resemble the findings of the current and various prior studies with faces: the early onset, the highest decoding accuracies between 400 and 600 ms, which reflect the Sustained Familiarity Effect, and the prolonged nature of familiarity processing, lasting well after 1000 ms (Dobs et al. 2019; Wiese et al. 2019; Ambrus et al. 2021). Expanding on the previous studies, the present research has demonstrated that highly personally familiar scenes exhibit a similar familiarity effect as highly familiar faces in terms of time course and scalp distribution of the neural signal. The sustained nature of this effect is consistent with an extended network of brain regions showing sensitivity to familiarity for natural videos in a recent fMRI study (Bainbridge and Baker 2022). One crucial difference between the familiarity classification dynamics of faces and scenes emerged in the earliest processing phase. While the familiarity effect for faces is characterized by an initial sharp increase shortly prior to 200 ms and an early peak before 300 ms, a similar effect for scenes is delayed and attenuated. The onset of the scene familiarity effect occurs only after 200 ms and it increases more gradually until it reaches its peak around 400 ms. From 400 ms onward, the classification performance for both face and scene familiarity declines, with the slight difference that scene familiarity information appears to be less sustained. A combination of two factors can be offered to explain these differences. First, it has been reliably shown that the visual processing of faces is prioritized, compared to any other stimulus categories (Langton et al. 2008; Kaneshiro et al. 2015; Contini et al. 2017; Morrisey et al. 2019). Second, compared to faces, scene images are more variable in terms of complexity and the arrangement of unique and distinct elements. This may lead to more variation in the time required for initial processing, decreasing signal-to-noise ratio in the early part of the signal. Overall, it is reasonable to assume a similar processing time course for familiarity in any other object category in future studies. While the very early emergence and secondary peak might be characteristic for only a few object categories (faces and very familiar objects; Ambrus, 2022), a general familiarity effect for any object category might look like the following: emergence around 300 ms, peak and plateau between 400 and 600 ms, and then a decline until the effect vanished at around 1000 ms. One can hypothesize that depending on the salience of the object category, a more sustained effect is also possible.

### Familiarity enhances category representations

While deeper, more sustained processing for familiar exemplars of an object category has been demonstrated thoroughly, only a few studies have looked at the specific corresponding processing benefits. Dobs and colleagues (2019) found that identity and gender information were modulated by familiarity even prior to 200 ms, suggesting an inherent feed-forward enhancement of familiar stimuli. More recently, Kovács et al. (2023) found that the that familiarity enhances identity decoding in the time range of 200-400 ms, while earlier neural representations appear to be relatively insensitive to the level of familiarity. Our study builds upon and extends these findings by demonstrating that familiarity enhances not only face representations but also those of non-face stimulus categories. The difference of category classification accuracies for familiar and unfamiliar stimuli revealed that familiarity enhances category classification performance both in an early (ca. 220 – 300 ms) and in a later (>400 ms) processing stage. The temporal dynamics of familiarity and category processing follow a similar pattern. An initial sharp increase is paired with a high peak early in the time course that is consistent with the N170, face-object discrimination, and scene (Jacques and Rossion 2006; Kaiser et al. 2020). This is followed by a plateau or drop in classification accuracy preceding a second, more gradual increase, leading to a more sustained effect with a late, secondary peak (between 400 and 600 ms). It is suggested that the two (early vs. late) phases in classification accuracy index different levels of processing within the processing hierarchy. The initial rapid feedforward signal is driven by low-level stimulus properties, which is later modified by more elaborate recurrent feedback signals (Contini et al. 2017). Our study found that the effects of stimulus category emerged earlier than familiarity signals, with category decoding being unaffected by familiarity until approximately 200 ms, in line with previous ERP findings on the onset of familiarity. This was followed by a decline in the accuracy of decoding for both familiar and unfamiliar stimuli. However, this decline was less pronounced for familiar stimuli, resulting in a more persistent category effect around 300 ms for familiar compared to unfamiliar stimuli. Finally, between 400 and 600 ms and beyond, category classification performance increases again. The rise in classification accuracy was more prominent for familiar stimuli, especially in the posterior regions of interest. Previous research (e.g., van de Nieuwenhuijzen et al. 2013; Kaneshiro et al. 2015), has shown that human faces have a significant impact on visual category classification analyses in terms of both the timing and strength of the effects. The results of our category decoding analysis suggest that familiarity enhances the efficiency of neural coding for both faces and scenes, making them easier to read out by classifiers and downstream brain regions. This indicates that familiarity is not only important for recognizing individual entities, but also contributes to more efficient perceptual and memory representations. Future studies could explore whether this effect extends to other types of stimuli beyond faces and scenes, particularly when human faces are not present in the stimulus categories.

## Limitations and future research directions

Although our results strongly support the generalizability, category- and stimulus-independence of the observed familiarity signal, here we have only compared familiarity between personally familiar faces and scenes. Both share some processing features, such as holistic representations (Konkle et al. 2010; Richler and Gauthier 2014; Kaiser and Cichy 2021) and appear to activate many similar occipito-temporal brain areas when highly familiar. One challenge that needs to be addressed in future studies is how the familiarity representation is implemented in visual cortex despite the separation of face and scene processing into dissociable category-selective processing streams (Kanwisher et al. 1997; Epstein and Kanwisher 1998; Taylor et al. 2007). One possibility is that familiarity is coded in a similar fashion across category-selective populations that gives rise to similar activation patterns in the EEG (e.g., via similar anterior to posterior gradients and focal processing areas), despite differences in spatial coding. Another possibility is that the effect is mediated by regions placed downstream from these category-selective visual areas, such as anterior and medial temporal regions, at the interface between perception and memory (Quiroga et al. 2005; Steel et al. 2021; Treder et al. 2021). These alternatives need to be explored in future fMRI studies. Furthermore, complete generalizability would require the comparison of several other non-face object categories and familiar stimuli from other modalities, such as auditory stimuli as well (Ambrus, 2022). Future studies should systemically investigate familiarity processing in a variation of non-face stimulus categories in the visual domain, to corroborate evidence from our study as well. Our results are further based on only highly personally familiar stimuli. Different degrees of familiarity (Li et al., 2022; Kovács et al, 2023), like perceptual and contextual familiarity (Kovács, 2020), should be related to the same processing mechanism but evoke slightly differential activations (Ambrus et al., 2021; Dalski et al., 2022a). Indeed, Beldzik and colleagues (2021)have shown that personally familiar scenes and known, but personally unfamiliar scenes, engage different brain networks related to an egocentric or more allocentric worldview, respectively. As the level of familiarity is closely related to and enhances face identity processing gradually (Dobs et al., 2019; Kovács et al (accepted)), it would be interesting to test if a similar identity effect exists for scenes as well, e.g., a differentiation between ones’ very well known, own bathroom and the bathroom at a railway station which is visited only irregularly. This, however, should only be a byproduct of scene processing, since identification is not the primary goal of scene perception and neural mechanisms should not be tuned explicitly towards it (Steel et al., 2021). Still, the results might shed more light on identity processing in general or how the brain distinguished between highly similar exemplars of the same stimulus category. To gain a comprehensive understanding of familiarity and its effect on neural processing, future studies need to vary analysis approaches and recording methods. This includes the univariate analysis of ERPs (Wiese et al., 2019) as well as the application of time-frequency analysis (Sáringer et al., 2023). Whereas right now, we were only able to speculate on the existence of a scene familiarity network, fMRI studies could examine brain areas related to scene familiarity processing for different familiarity qualities.

Summary

The present study examined the generalizability of familiarity information between two different stimulus categories (faces and scenes). Previous face familiarity results, related to the established dynamics of face familiarity processing as well as the generalizability of this familiarity signal across participants, were confirmed. This familiarity classification was extended to the stimulus category of scenes. The temporal dynamics of scene familiarity revealed an onset shortly around 200 ms and a peak around 400 ms, followed by a slow decrease. Besides an earlier onset and a sharper rise of face familiarity information the familiarity signal generalized in a remarkably robust way between faces and scenes emerged. We further found good generalizability of the familiarity signal across stimulus categories not only within-participant but also cross-participant, indicating that the previously described familiarity effects are not face-specific. Our results thus emphasize the robust and sustained nature of a general familiarity effect, independent of stimulus category and participant. Finally, we tested whether and how familiarity modulates the neural dynamics of category representations. We found that familiarity enhances the category information in both early and late processing stages, leading to a deeper processing of familiar stimuli.

## Data and code availability

Data and code are uploaded to OSF (https://osf.io/m9q74/). We have not uploaded the experimental stimuli, as the conditions of our ethics approval do not permit public archiving of these personal images. The entire data and stimulus sets will be made available to interested researchers following completion of a data sharing agreement and approval by the local ethics committee. No part of the study procedures or analyses was pre-registered prior to the research being conducted.

## Ethics statement

The study was conducted in accordance with the guidelines of the Declaration of Helsinki and was approved by the ethics committee of the Friedrich-Schiller-Universität Jena.

## Acknowledgments

D.K. is supported by the Deutsche Forschungsgemeinschaft (DFG; SFB/TRR135 – “Cardinal Mechanisms of Perception”, Project Number 222641018), a European Research Council (ERC) starting grant (ERC-2022-STG 101076057), and “The Adaptive Mind”, funded by the Excellence Program of the Hessian Ministry of Higher Education, Science, Research and Art.

## Author contributions

GK, GGA and HK designed the experiment. HK wrote the experimental code and performed the data collection. GGA analyzed the data. DK and RS provided consultation on the methods and analysis. HK, DK, RS, GK, and GGA wrote the manuscript.

## Declaration of interests

The authors have no competing interests to declare.

## Supplementary Materials

**Supplementary information are available under https://osf.io/m9q74/**

**Supplementary 1**. Statistics for the within-subject and cross-participant time-resolved classification analyses. All sensors and pre-defined regions of interest

**Supplementary 2**. Statistics for the within-subject and cross-participant temporal generalization analyses. All sensors and pre-defined regions of interest

**Supplementary 3**. Statistics for the within-subject and cross-participant spatiotemporal searchlight analyses

## Notes

https://osf.io/m9q74/

### Competing Interest Statement

The authors have declared no competing interest.

