## Supplementary Data for "Your place or mine? The neural dynamics of personally familiar scene recognition suggests category independent familiarity encoding"

**Supplementary Information 1**

In the **time-resolved classification** procedure, linear discriminant analysis classifiers were iteratively trained at each of the 140 time points (ranging from -200 to 1200 ms) to distinguish between the classes of interest (either familiar versus unfamiliar stimuli in the familiarity classification analysis, or faces versus scenes in the category classification analysis). These classifiers were then used to evaluate the accuracy of decoding either for a portion of the data from each participant individually (in the within-participant analyses) or for data from a participant who was not included in the training set (in the leave-one-participant-out analyses).

**Temporal generalization** analyses were conducted in a similar fashion. Here, classifiers trained at a given time-point were used to test held-out data form all other time points. This results in a cross-temporal (training-times × testing-times) classification accuracy matrix where similar information-processing is indicated where classifiers successfully generalize from one time point to another (King & Dehaene, 2014).

In the **spatio-temporal searchlight analysis**, we systematically tested each channel by training and testing on data from that channel and its adjacent electrodes. I.e., for each channel and its neighboring electrodes, we conducted a time-resolved analysis in the manner described above.

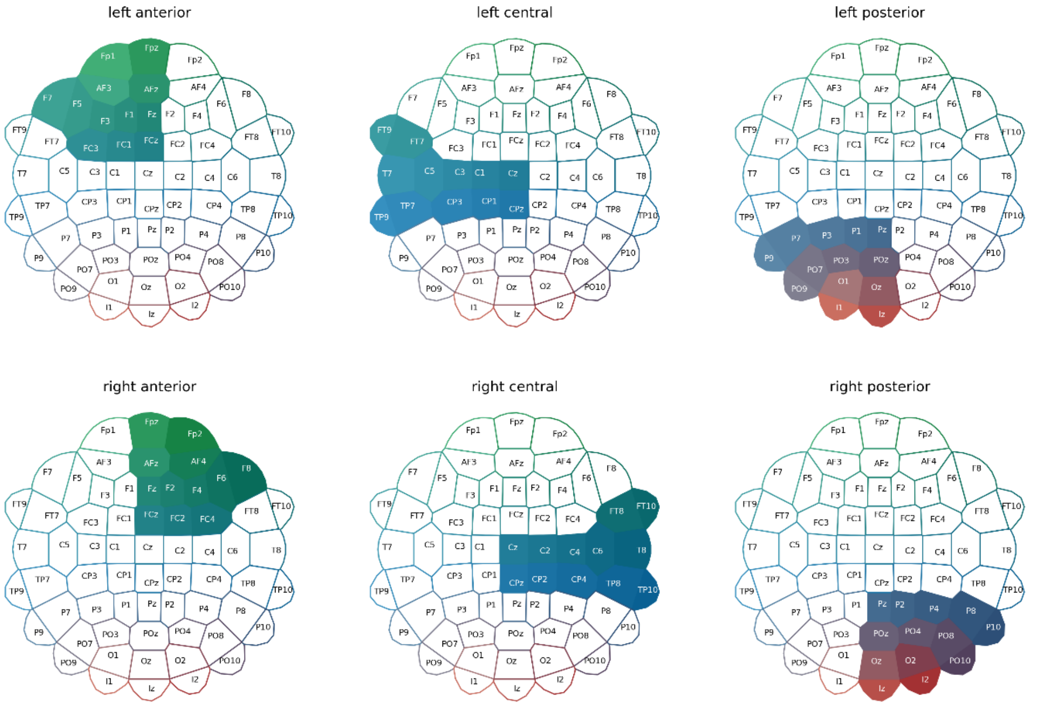


**Supplementary Figure S1. Regions of interest.** Six scalp locations along the medial (left and right) and coronal (anterior, center, and posterior) planes were defined. Supplements the Methods section in the main text.

Time-resolved classification and temporal generalization analyses were performed on all channels and pre-defined regions of interest. These ROIs were based on Ambrus et al. (2022; 2019, 2021) and Dalski et al. (2022). Six scalp locations along the median (left and right) and coronal (anterior, center, and posterior) planes (see **Figure S1**) were defined and used for training and testing in separate analyses.

**Supplementary Information 2**

**Within-participant familiarity classification.** To probe the temporal dynamics of familiarity representation for faces and scenes on the participant level, within-subject classification analyses were conducted. To balance the trial counts, ERPs for all four personally familiar identities were used, while dropping the trials for one of the five unfamiliar identities (preferably of the gender that was overrepresented in the familiar faces stimulus set). To balance the familiar scene data, an equal number of familiar and unfamiliar images were retained.

In within-participant familiarity classification, for faces, trials for each identity were iteratively held out for testing. As all images for scenes all depicted the same locations (i.e., the apartment of the participant and an unfamiliar apartment), the scene stimulus set was divided into chunks of equal size (containing 4-4 images, to match the identity division in faces) for the purposes of train-test splits. The resulting accuracy scores for all train-test splits were then averaged.

**Results**

**Within-category classification** over all electrodes (**Figure S2A**) revealed strong significant clusters (cluster *p*s < 0.0001), for faces starting at 170 ms, with a peak at 410 ms (peak Cohen's *d* = 1.331), and for scenes starting at 180 ms, with a peak at 720 ms (peak Cohen's *d* = 1.333), both lasting to the end of the epoch. Spatio-temporal searchlight analyses revealed similar patterns, both for faces and scenes, robust clusters (cluster *p*s < 0.0001), encompassing all electrodes, have been observed for faces (onset: 130 ms, peak at 460 ms over PO8, peak Cohen's *d* = 1.890) as well as scenes (onset: 80 ms, peak at 450 ms over POz, peak Cohen's *d* = 1.535). Temporal generalization results (**Figure S4A-B**) yielded single, sustained, rectangular clusters both in the case of faces (between 130 to 1190 ms train times and 10 to 1190 ms test times, cluster p < 0.0001, peak Cohen's d = 1.515) and scenes (between 80 ms to the end of the epoch for train times and 160 ms to the end of the epoch for test times, cluster p < 0.0001, peak Cohen's d = 1.333). Results of the time-resolved within-subject classification and temporal generalization analyses for the pre-defined regions of interest can be found in **Supplementary Table 1A-C** and **Supplementary Table 2A-B.** For the details of the searchlight results, see **Supplementary Table 3A-B**.

**Cross-category classification** over all electrodes (**Figure S2B**) yielded strong significant clusters (cluster *p*s < 0.0016), for training on scenes and testing on faces between 240 and 770 ms, with a peak at 340 ms (peak Cohen's *d* = 1.176), and for training on faces and testing on scenes between 230 and 650 ms, with a peak at 420 ms (peak Cohen's *d* = 1.047). Again, spatio-temporal searchlight analyses revealed similar patterns, in both directions, robust clusters (cluster *p*s < 0.0001), encompassing all electrodes, have been observed; for scenes to faces (onset: 110 ms, peak at 440 ms over TP8, peak Cohen's *d* = 1.890) as well as faces to scenes (onset: 120 ms, peak at 430 ms overTP10, peak Cohen's d = 1.425). In both directions, temporal generalization (**Figure S4C-D**) yielded single, sustained, rectangular clusters, for scenes to faces (120 ms to the end of the epoch for train times, 50 ms to the end of the epoch for test times, cluster *p* = 0.0014, peak Cohen's *d* = 1.167) and for faces to scenes (80 ms to the end of the epoch for train times, 140 ms to the end of the epoch for test times, cluster *p* = 0.0073, peak Cohen's *d* = 1.019). Results of the time-resolved within-subject cross-classification and temporal generalization analyses for the pre-defined regions of interest can be found in **Supplementary Table 1D-F** and **Supplementary Table 2C-D.** For the details of the searchlight results, see **Supplementary Table 3C-D**.

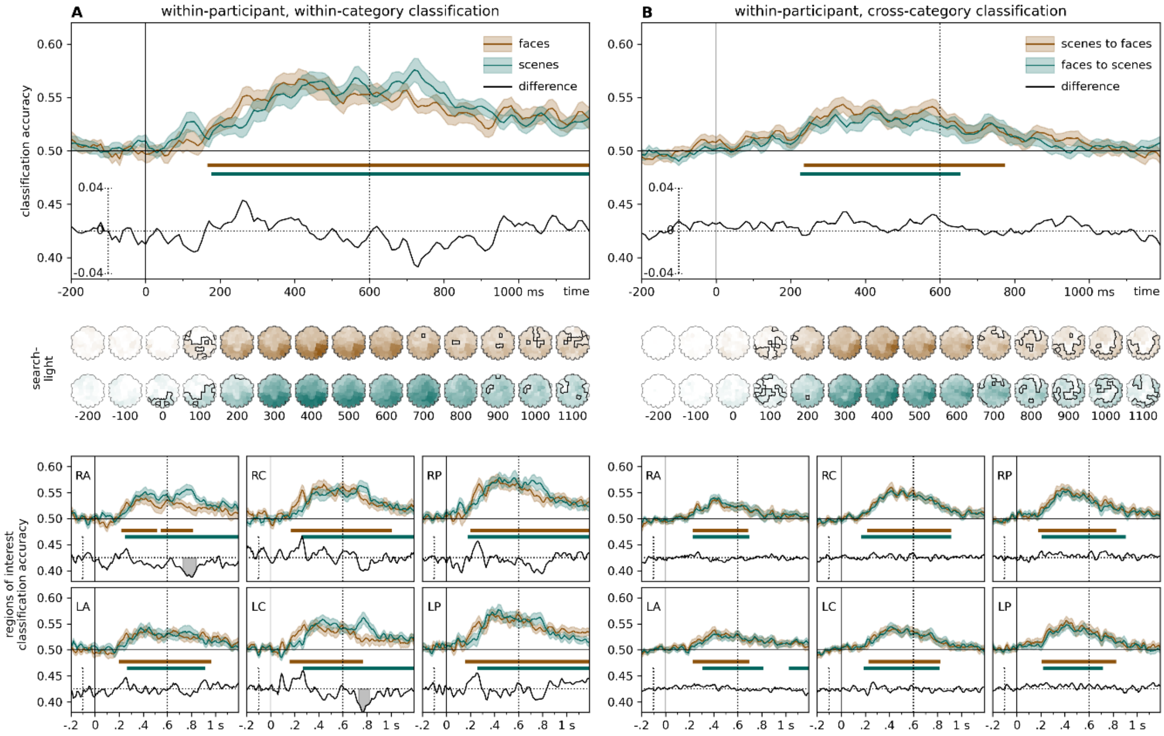


**Supplementary Figure S2. Time-resolved, within-participant classification.** Classifiers were trained to categorize ERPs for familiar and unfamiliar stimuli. **(A) within-category classification**. Training and testing for familiarity was performed within the same stimulus category. **(B) cross-category classification.** Training and testing for familiarity was performed on different stimulus categories. Two-sided cluster permutation tests, $p < 0.05$.

Spatio-temporal searchlight results are shown as scalp maps, with classification accuracy scores averaged in 100 ms steps. Sensors and time points belonging to the significant cluster when tested on faces are shown in the top row, sensors and time points belonging to the significant cluster when tested on scenes are shown in the bottom row. (Two-sided spatio-temporal cluster permutation tests, $p < 0.05$).

RA/LA: right/left anterior, RC/LC: right/left central, RP/LP: right/left posterior. For detailed statistics, see **Supplementary Table 1A-F**, and **Supplementary Table 3A-D**.

**Supplementary Information 3**

**Within-participant stimulus category classification.** To investigate the neural dynamics of stimulus category representations in relation to familiarity at the participant level, we conducted separate within-subject face versus scene classification analyses on evoked responses for familiar and unfamiliar trials. The train-test selection process followed similar procedures as described for the within-participant familiarity classification analysis.

**Results**

Time-resolved leave-one-participant-out analyses (**Figure S3**) for both familiar and unfamiliar stimuli revealed strong (cluster *p*s < 0.0001), sustained clusters, with an early onset. For familiar stimuli, the cluster onset was at 90 ms, with a peak at 160 ms (peak Cohen's *d* = 3.350), while for unfamiliar stimuli, the onset of the significant cluster was at 100 ms, with a similar peak at 160 ms (peak Cohen's *d* = 3.286). A significantly different late cluster (between 480 and 670 ms, peak at 530 ms, cluster p = 0.0042, peak Cohen's *d* = 0.710) was observed for stimulus category decoding between familiar and unfamiliar stimuli, with higher classification accuracies seen for familiar stimuli (see **Figure S3**). Searchlight analyses revealed robust stimulus category classification effects both for familiar and unfamiliar stimuli (cluster *p*s < 0.0001), both peaking at 160 ms over the PO8 electrode region (peak Cohens *d*s = 4.898 and 4.345, respectively).

Temporal generalization results (**Figure S4E-F**) yielded robust (cluster *p*s < 0.0001), sustained, rectangular clusters both in the cases of familiar (from 90 ms train time and 90 ms test time, peak Cohen's *d* = 3.126) and unfamiliar stimuli (from 100 ms train time and 100 ms test time, peak Cohen's *d* = 3.286).

Results of the within-participant stimulus category time-resolved cross-classification and temporal generalization analyses for the pre-defined regions of interest can be found in **Supplementary Table 1G-I** and **Supplementary Table 2E-F.** For the details of the searchlight results, see **Supplementary Table 3E-F**.

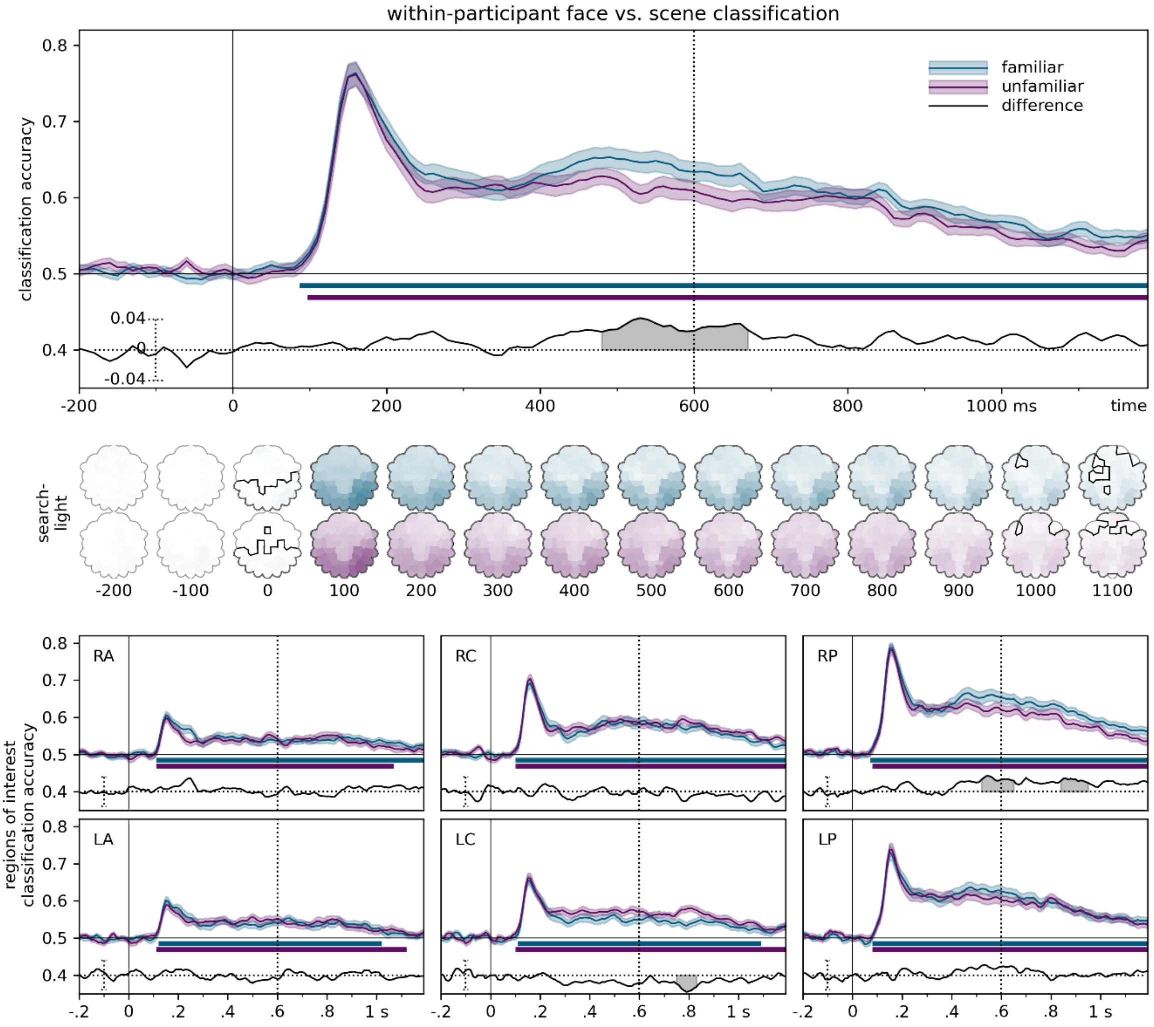


**Supplementary Figure S3. Time-resolved, within-participant classification of stimulus category.** Classifiers were trained, within-participant, to categorize ERPs stimulus category (faces vs. scene), for familiar and unfamiliar trials separately. Two-sided cluster permutation tests, $p < 0.05$. Spatio-temporal searchlight results are shown as scalp maps, with classification accuracy scores averaged in 100 ms steps. Sensors and time points belonging to the significant cluster when tested on familiar stimuli are shown in the top row, sensors and time points belonging to the significant cluster when tested on unfamiliar stimuli are shown in the bottom row. (Two-sided spatio-temporal cluster permutation tests, $p < 0.05$). The vertical line at 600 ms denotes the end of the stimulus presentation. RA/LA: right/left anterior, RC/LC: right/left central, RP/LP: right/left posterior. For detailed statistics, see **Supplementary Table 1F-I.**, and **Supplementary Table 3E-F**.

**Supplementary Information 4**

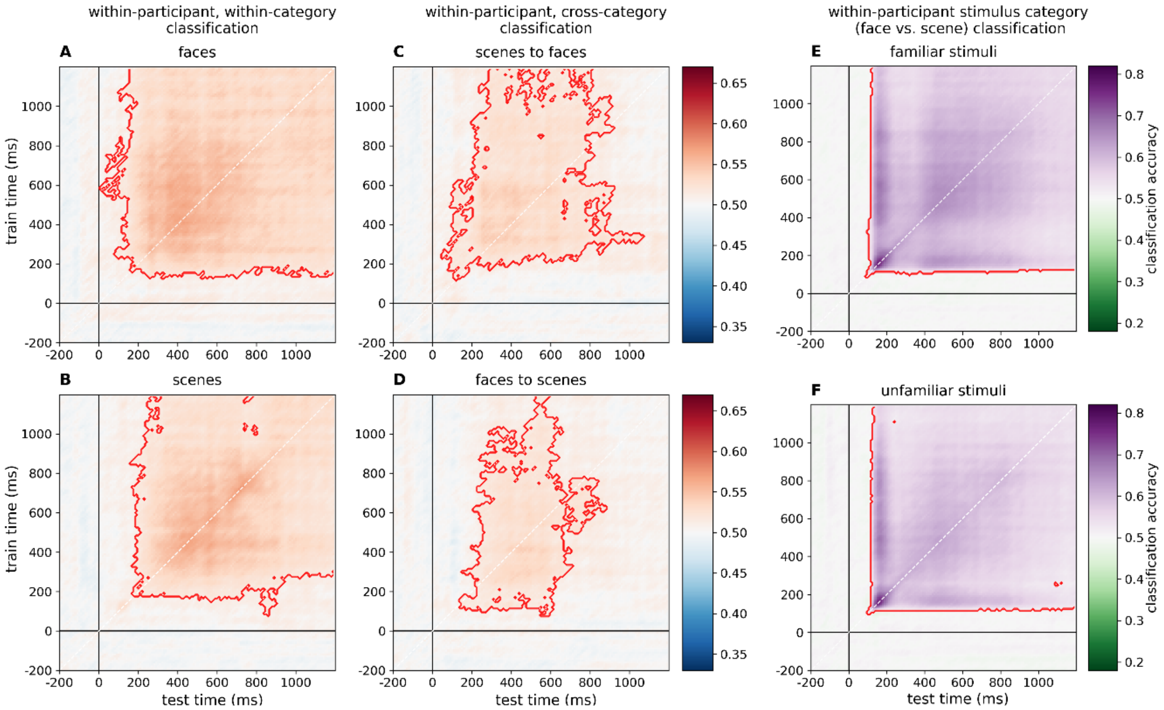


**Supplementary Figure S4. Within-participant temporal generalization analyses** of familiarity and category. **(A, B) within-category classification.** Classifiers were trained to categorize ERPs for familiar and unfamiliar stimuli. Training and testing for familiarity was performed within the same stimulus category. **(C, D) cross-category classification of familiarity**. Training and testing for familiarity was performed on different stimulus categories. **(E, F) cross-category classification of stimulus category.** Classifiers were trained, within-participant, to categorize ERPs stimulus category (faces vs. scene), for familiar and unfamiliar trials separately. Two-sided cluster permutation tests, $p < 0.05$. For the results of the temporal generalization analyses in the pre-defined regions of interest, see **Supplementary Table 2.**
